# Investigating the mode of action for wasting produced by tetrachlorodibenzo-p-dioxin (TCDD) in rats using transcriptomics: Evidence for roles of AHR and ARNT in circadian cycling

**DOI:** 10.1101/2024.03.02.583130

**Authors:** Melvin E. Andersen, A. Rasim Barutcu, Michael B. Black, Joshua A. Harrill

## Abstract

Single, high doses of TCDD in rats caused wasting, a progressive loss of 30 to 50% body weight and death within several weeks. To identify pathway perturbations at or near doses causing wasting, we examined differentially gene expression (DGE) and pathway enrichment in centrilobular (CL) and periportal (PP) regions of female rat livers following 6 dose levels of TCDD – 0, 3, 22, 100, 300, and 1000 ng/kg/day, 5 days/week for 4 weeks. At the higher doses, rats lost weight, had increased liver/body weight ratios and nearly complete cessation of liver cell proliferation, signs consistent with wasting. DGE curves were left shifted for the CL versus the PP regions. Canonical Phase I and Phase II genes were maximally increased at lower doses and remained elevated at all doses. At lower doses, ≤ 22 ng/kg/day in the CL and ≤ 100 ng/kg/day, upregulated genes showed transcription factor (TF) enrichment for AHR and ARNT. At the mid- and hi-dose doses, there was a large number of downregulated genes and pathway enrichment for DEGs showed downregulation of many cellular metabolism processes including those for steroids, fatty acid metabolism, pyruvate metabolism and citric acid cycle. There was significant TF enrichment of the hi-dose downregulated genes for RXR, ESR1, LXR, PPARalpha. At the highest dose, there was also pathway enrichment with upregulated genes for extracellular matrix organization, collagen formation, hemostasis and innate immune system. TCDD demonstrates most of its effects through binding the aryl hydrocarbon receptor (AHR) while the downregulation of metabolism genes at higher TCDD doses is known to be independent of AHR binding to DREs. Based on our results with DEG, we provide a hypothesis for wasting in which high doses of TCDD shifts circadian processes away from the resting state leading to greatly reduced synthesis of steroids and complex lipids needed for cell growth and producing gene expression signals consistent with an epithelial-to-mesenchymal transition in hepatocytes.

## Introduction

Individuals involved in production of 2,4,5-trichlorophenol (2,4,5-T) developed chloracne, porphyria cutanea tarda, anorexia and weight loss (Poland *et al*., 1971; Girer *et al*., 2020). TCDD was found as a contaminant in 2,4,5-T production and, in toxicology studies, high single doses of TCDD (50 or 100 µg/kg) in rats caused anorexia, wasting (loss of up to 50% body weight) and death within two to three weeks (Gasiewicz *et al*., 1980). Pair-fed control animals also lost weight and died at a similar time after dosing (Christian *et al*., 1986). Hepatocytes of treated rats were enlarged compared to pair-fed controls and contained nuclei that varied in size and number. When TCDD-treated and control rats were fed by total parenteral nutrition (TPN), where the intravenous (IV)-administered nutrition is the only source of nutrition (Gasiewicz *et al*., 1980), TCDD-treated rats still died within several weeks while TPN-controls were not affected. Histologically, the livers of TCDD treated TPN-rats weighed 2 to 3-times those of TPN-fed control rats and were severely necrotic, swollen, and disorganized. While reduced nutritional intake was consistent with lethality, the alterations in liver architecture with TPN supplementation indicated that TCDD caused severe disruption of metabolic processing of nutrients. Treateatment of mice with an antibody to tumor necrosis factor reduced mortality from TCDD (Taylor *et al*., 1992), but these early observations on TNF were not pursued. Importantly, the observation of wasting in rats and mice and anorexia and weight loss in humans with TCDD (Poland *et al*., 1971; Gasiewicz *et al*., 1980) indicate that rodents and primate species have similarities in responses to high doses of TCDD.

TCDD is an agonist for AHR, a β-helix loop helix (β-HLH) transcription factor (McIntosh *et al*., 2010). This transcription factor binds TCDD with high affinity and the liganded protein interacts with another β-HLH protein, ARNT (AHR nuclear translocator also known as hypoxia inducing factor – 1β (HIF1β). This molecular complex, consisting of AhR, ARNT and TCDD, along with other co-factors, binds to dioxin-response element (DRE) promoters to activate transcription of various genes, including *Cyp1a1, Cyp1a2, Cyp1b1* and *Nqo1* (Dere *et al*., 2011). At higher doses, TCDD also caused downregulation of genes involved in cholesterol and fatty acid metabolism in mice (Sato *et al*., 2008), a response that did not require AhR binding to canonical DREs that are present in the differentially expressed Phase I and II genes (Tanos *et al*., 2012). Both AHR and ARNT have biological activity in the absence of activating ligands such as TCDD. ARNT forms a heterodimer with HIF1A, and this dimer regulates transcription of hundreds of genes in response to reduced oxygen levels (Prabhakar, 2020). Similarly, unliganded AHR has other biological functions, binding with CDK4 and Retinoblastoma (RB) to produce a cellular state permissive for cell proliferation (Barhoover *et al*., 2010) and liganded AHR inhibits invasiveness and metastatic features of breast cancer cells (Hall *et al*., 2010). AHR also has effects on circadian rhythms (Jaeger and Tischkau, 2016; Jaeger *et al*., 2017; Tischkau, 2020). AHR-KO mice have altered circadian rhythmicity and increased amplitude of circadian responsive genes in peripheral tissues. TCDD at a dose of 30 ug/kg once every 4 days (7 total doses) abolished circadian regulation of hepatic metabolic activity in mice (Fader *et al*., 2019).

A decade ago, studies were initiated to understand the role of AHR in liver responses in female rats using both wild-type and AHR-knock out (AHR-KO) female Sprague-Dawley rats. Female rats were used in the original study because liver responses, especially cancer, in this sex and strain of rats were being used as the basis for cancer risk assessments. Two reports of this work have appeared – one describing phenotypes for liver and kidney in AHR-KO rats (Harrill *et al*., 2013) and another showing that the AHR-KO rats were insensitive to repeat exposures of TCDD (Harrill et al., 2016). In the 1000 ng/kg/day group, there were decreases in body weight and increases in liver/body weight ratios, indicating that these doses were nearing conditions consistent with wasting.

In this study, we sought to gain insight into the metabolic mode of action (MOA) for wasting by examining DGE in periportal (PP) and centrilobular (CL) regions of livers from the rats in the earler stdy receiving various doses of TCDD (0, 3, 22, 100, 300 and 1000 ng/kg/day) for 4 or 5 days/week for 4 weeks for a total of 19 doses. This dataset provided an opportunity to examine liver gene expression changes occurring at doses at or very near those associated with wasting (Harrill *et al*., 2016). Our analysis examined the dose response for the number of DEGs, TF enrichment for DEGs and evaluated pathway enrichment for the DEGs at various doses. With TCDD at the higher dose, there was significant downregulation of many metabolism pathways, including lipid and steroid metabolism, fatty acid oxidation and biological oxidations. We integrate our results with the known biology affected by TCDD to offer a hypothesis for the mode of action (MOA) of TCDD in causing wasting-like responses. The hypothesis proposes that AhR and A have central roles in circadian cycling in peripheral tissues that are disrupted by the presence of a persistent, poorly metabolized AhR ligand (TCDD) that sequesters AHR and ARNT as a dimer and limits their ability to interact with other TF partners. In addition, persistent upregulation of CYPs (1A1, 1A2, and 1B1) cause enhanced clearance of various small molecules such as melatonin and alters control of core components of circadian circuitry. Longer chain perfluorinated fatty acids (PFAS) also cause wasting-like toxic signs in rodents and non-human-primates. These compounds also cause profound down-regulation of metabolism pathways for lipogenesis in human hepatocytes treated in culture for 14-days (Barutcu *et al*., 2024).

## Materials and Methods

### Animal Care, Husbandry and Treatment

The Hamner Institutes for Health Sciences Laboratory Animal Resources and Technical Support Facility was fully accredited by the Association for Assessment and Accreditation of Laboratory Animal Care during the experiment. All breeding and experimental procedures were approved by the Hamner Institutes’ Institutional Animal Care and Use Committee.

Wild-type and AHR-KO rats harboring a 29-bp deletion mutation in exon 2 of the AHR gene were used in this study. Wild-type and AHR-KO rats were generated from heterozygous breeding stocks maintained on a Sprague-Dawley outbred background and bred at the Hamner Institutes for Health Sciences, Research Triangle Park, North Carolina (Harrill *et al*., 2013). Animal care, husbandry and treatment of wild-type and AHR-KO rats are described (Harrill *et al*., 2016). Briefly, after weaning female wild-type and AHR-KO rats of like genotypes were housed two per cage in solid-bottom polycarbonate microisolator cages containing Alpha-dri cellulose bedding (Shepherd Specialty Papers, Kalamazoo, MI) with free access to NIH-07 certified feed (Zeigler Brothers, Gardners, PA) and reverse osmosis purified water (Hydro Systems, Durham, NC). All breeding stocks and test animals were housed in rooms maintained between 20 and 26 °C with 40 to 60% relative humidity and standard 12 hour light:dark cycle (0700-1900).

At 49 ± 4 days of age, one week prior to the start of dosing (56 ± 4 days of age), rats were weighed, randomized into treatment groups, and housed as previously described. Rats with body weights greater or less than 20% of the overall mean body weight for the entire test population were not included in the study. The study was conducted using two equivalently sized blocks of animals (n = 5 rats / dose group / genotype / block) which were derived from consecutive litters born to the breeding stocks. Transcriptomic data was combined across the two blocks (n = 10 rats / dose group / genotype).

Rats were dosed by oral gavage with varying concentrations of TCDD (0, 3, 22, 100, 300, 1000 ng/kg/day), 4-5 days / week for four weeks; a total of 19 doses. Corn oil was used as the test vehicle with a dose volume of 5 mL/kg body weight. Dose solution preparation and confirmation of TCDD concentrations in dose solutions by gas chromatography-low resolution mass spectrometry (GC/LRMS) was as described (Harrill *et al*., 2016).

### Sample Collection and Preservation

At necropsy, rats were deeply anesthetized with CO_2_ and exsanguinated by transection of the abdominal aorta. The liver was removed and sections from the middle of the left liver lobe (1 mm thick, < 10 mm long) were collected, covered in Tissue-Tek® Optimal Cutting Temperature (OCT) compound (VWR International, Padnor, PA) and frozen on a stainless-steel plate placed on a bed of dry ice. Frozen liver sections were stored at -80°C prior to sectioning for laser capture microdissection.

### BrdU and Ki67 immunohistochemical analysis

Paraffin embedded rat livers from WT and AhR KO rats treated with TCDD (0, 3, 22, 100, 300, 1000ng/kg/day for 28 days, n=5 per group) were sectioned, and 5 µm sections from each block were stained for the presence of either Ki67 or BrdU using primary antibodies from Santa Cruz Biotechnology. Hepatocytes that incorporated Ki67 or BrdU were identified by brown to black pigment within their nuclei. Sections of small intestine served as positive controls. Both Ki67 and BrdU labeled, as well as unlabeled hepatocellular nuclei, were recorded from a minimum of five 400x fields (40x lens x 10x ocular) per section of liver. A total of 1000 hepatocytes were scored per section. The labeling index was expressed as a percentage of the number of labeled nuclei counted (hepatocellular) per total number of nuclei (hepatocellular).

### Laser Capture Microdissection (LCM)

Frozen tissue blocks from livers mounted on a brass chuck and placed into a cryostat equilibrated to -14°C. Frozen sections 12 µm in thickness were cut and mounted on MMI RNase-free MembraneSlides (Molecular Machines & Industries, Eching, Germany). Between three and five 12 µm sections from each tissue block were placed on a slide. Freshly-cut slides were allowed to air dry for 10 seconds in the cryostat chamber then fixed in ice-cold 70% ethanol for 3 min, rinsed in RNase free water for 30 s and stained for 15 s in a 0.22 µM filtered hematoxylin solution. Slides were then quickly rinsed twice with RNAse-free water, dehydrated by 70% followed by 100% ethanol (15 s each) and allowed to air dry at room temperature for 3 min. LCM was then performed on each slide immediately following the hematoxylin quick-staining protocol. The described quick-staining protocol was adapted from previously published methods designed to preserve the integrity of biological macromolecules (Stemmer and Dietrich 2011).

An MMI CellCut Plus LCM instrument with MMICellTools software (v3.47) was used to collect PP and CL hepatocytes for transcriptomic analysis. Stained slides were placed on the LCM dissecting stage and liver sections were visually surveyed using a 4X objective. Approximately 200 vascular structures (i.e., portal veins or central veins) were marked with reference points across all liver sections present on the slide. A higher magnification objective was then used to review reference points, discriminate between portal and centrilobular vascular structures and plot “horseshoe” shaped areas for LCM and tissue collection Areas for dissection around PP reference points were traced until a total of 2×10^6^ µm^2^ was demarcated for dissection. Reference points were then sequentially dissected and collected using a MMI Diffuser Cap and the MMI CellCut Plus Caplift mechanism. Following collection of all PP reference points in the sample, the Diffuser Cap was removed, filled with 150 µL of β-mercaptoethanol supplemented TR1 buffer as supplied with the PAXgene Tissue RNA Kit (PreAnalytix GmbH, Hombrechtikon, Switzerland), inverted and stored at 4°C for 20 minutes. The dissection process was then repeated as described for CL reference points. Figure 1 shows representative hematoxylin quick-stained sections for periportal (A-B) and centrilobular (C-D) reference points from cryopreserved rat liver sections before (A,C) and after (B,D) laser capture microdissection. Sections were labeled using a hematoxylin quick-stain procedure.

**Figure 1:**
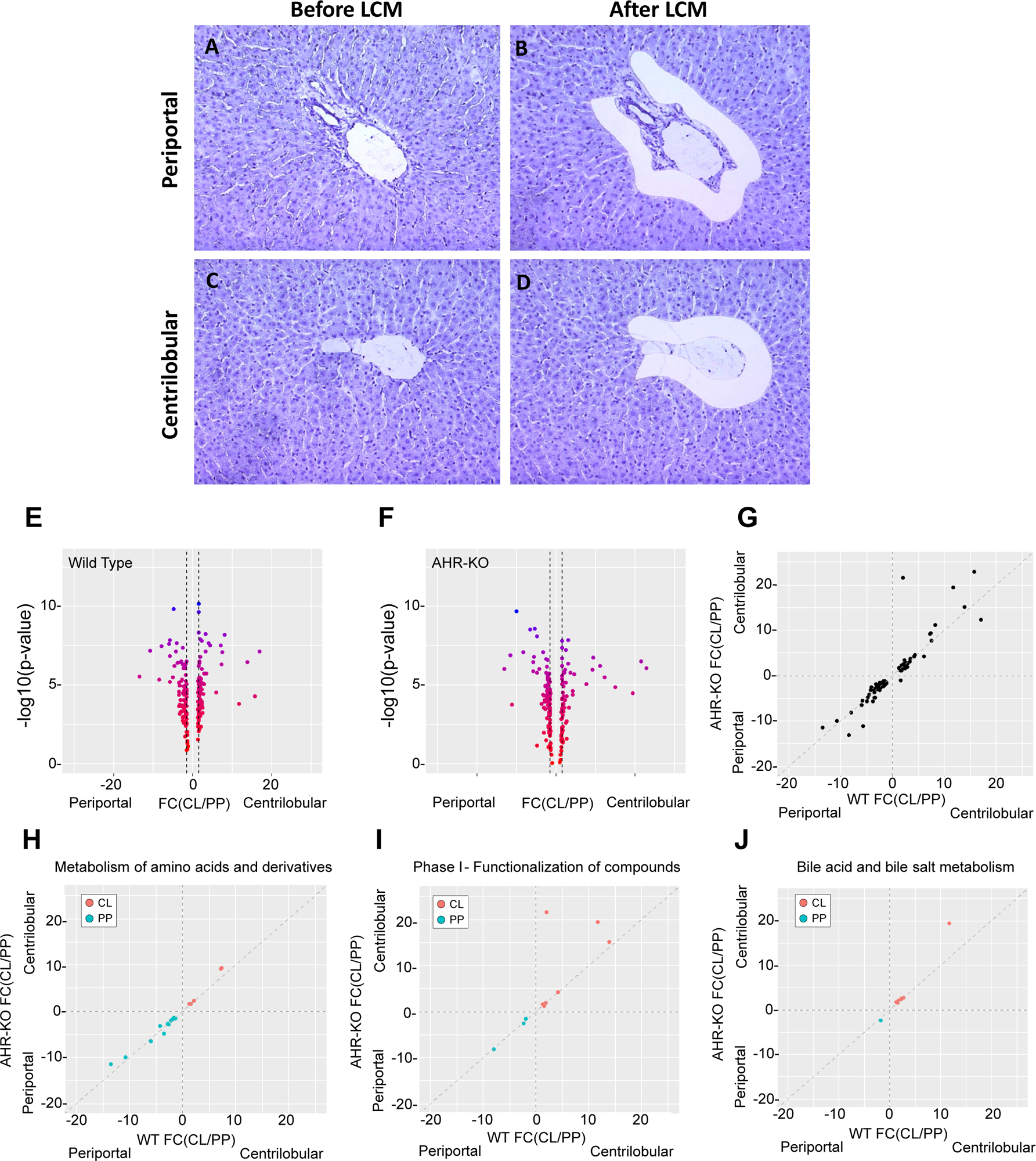
H&E staining showing morphology of CL and PP regions before and after laser capture microdissection. (A, B) PP region; (C and D) CL region. (E, F) Volcano plots of fold-change (FC) in expression between periportal and centrilobular hepatocytes in WT (E) and AHR-KO (F) vehicle control livers. Points to the left are expressed higher in PP hepatocytes while points to the right are expressed higher in CL hepatocytes. The color gradient flowing from red to blue indicates level of statistical significance. (G) Comparison of FC estimates across WT and AhR-KO rats. Points along the unity line have equivalent levels of zonal enrichment in WT and AhR-KO rats. Points distant from the unity line indicate that AHR status affects the zonal expression of these genes. Three of the genes with the greater expression in CL compared to PP region (Cyp1a2, Cyp7a1 and Cxl13) are marked in the figure. (H) Correlation plots for select Reactome pathways demonstrate that the DEGs in the metabolism of amino acids and derivative pathway are primarily expressed at higher levels in the PP hepatocytes while genes associated with Phase I functionalization of compounds (I) and bile acid/bile salt metabolism (J) are primarily expressed at higher levels in the centrilobular region. Gene expression results from which this figure was generated are in **TableS1**.

### RNA Extraction, Processing and Gene Expression Microarray Measurements

Total RNA was extracted from PP and CL hepatocytes using a PAXgene Tissue RNA Kit (PreAnalytix GmbH, Hombrechtikon, Switzerland). Total RNA was amplified, and cDNA prepared using the Ovation® Pico WTA System V2 (NuGEN Technologies, San Carlos, CA). The concentrations of amplified cDNA were quantified using a NanoDrop 2000 spectrophotometer and 5 µg of cDNA was labeled and fragmented using a NuGEN Encore® Biotin Module. For each sample, a hybridization cocktail was prepared, plated, and loaded into an Affymetrix GeneTitan system for hybridization (16 h) on an HT_Rat230_PM array plate (GPL16985), washing and scanning. All kits and procedures were performed according to manufacturers’ recommended protocols. The gene expression results have been deposited in the National Center for Biotechnology Information (NCBI) Gene Expression Omnibus (accession no.: GSE72506). Microarray data were processed using robust multi-array average normalization with a log2 transformation prior to the analysis steps described below.

### Analysis of Gene Expression in Vehicle Control Tissues

Affymetrix CEL file values were normalized by RMA and log2 transformed for analyses (Irizarry *et al*., 2003). The log2-transformed microarray data from vehicle treated PP and CL hepatocytes from WT and AHR-KO, respectively, were analyzed by one-way ANOVA to identify dose dependent changes in gene expression relative to vehicle controls, with orthogonal linear contrasts to test for differences at each individual treatment dose. Probability values were adjusted for multiple comparisons using a false discovery rate (FDR) of 5 % (Benjamini and Hochberg, 1995). Probes were considered significantly different in the two cell populations at FDR < 0.05 with an absolute value of fold change greater than or equal to 1.5-fold (|FC| > 1.5). Promiscuous or unannotated probe sets were removed the list of probes which passed the significance criteria and probes interrogating the same gene were aggregated to calculated gene level fold-change (FC).

### Pathway Enrichment for Differentially Expressed Genes

For each dose group, pathway enrichment was performed for both upregulated and downregulated genes in the PP and CL regions of the liver. For each treatment group, all genes with |FC|>1.5 and FDR<0.05 were collected to examine pathway enrichment using the Reactome ontology (Reactome.org) and for assessing transcription factor (TF) enrichment of DEGs using the 2022 ChEA database in Enrichr that allows reconstruction of a network of TFs based on shared overlapping targets and DNA-binding site proximity. The development and continuous enhancement of the Enrichr platform has been described in several publications (Chen *et al*., 2013; Kuleshov *et al*., 2016; Xie *et al*., 2021). The output from ChEA was collected and used to create tables for TFs associated with different treatments for different times of exposure. We also used Reactome pathway enrichment results with CytoScape 3.9.1 (Cytoscape.org) to generate GO-association network graphs, based on enrichment for up- and downregulated DEGs from the Reactome ontologies (https://reactome.org). Differentially expressed transcription factor (TF) genes were identified from TFCheckPoint.org and curated based on experimental evidence for (a) regulation of RNA polymerase II and (b) specific DNA binding activity.

## Results

### Region-specific Gene expression in vehicle control treated AHR-KO and AHR-WT rats

The process of laser capture microdissection was visualized by H&E staining of CL and PP hepatocytes before and after isolation of hepatocytes showing the pattern of harvesting from the two regions (**Figure 1 A-D**). Comparison of gene expression from these regions from WT vehicle controls (i.e., 0 ng TCDD/kg-day) identified 718 probes with differential expression between the two cell populations (**Table S2; Figure 1 E**). Comparable numbers of probes had higher expression in PP or CL hepatocytes, respectively, in this comparison with the highest differential zonal expression in the range of 15-20 fold. Similarly, comparison of gene expression data from PP and CL hepatocytes samples from AHR-KO vehicle controls identified 573 probes with differential expression (**Table S1**, **Figure 1F**). Again, comparable numbers of probes had higher expression in PP or CL hepatocytes, respectively, with the ratios of the highest zonal expression in the range of 15-20 fold.

Among the 699 genes that were significantly differentially expressed in CL and PP hepatocytes (|FC|>1.5; FDR<0.05) in the control treated rats (**Figure1 H-J)** were genes involved in the metabolism of amino acids (higher levels in the PP region) and genes associated with Phase I functionalization of compounds and bile acid/bile salt metabolism (higher in the CL region). These observations of different metabolic competence in the regions are consistent with zonal stratification of liver function (Jungermann, 1992; Jungermann and Kietzmann, 1996; Gebhardt and Matz-Soja, 2014; Halpern *et al*., 2017) and indicate that the sampling protocol used in this study successfully captured distinct populations of CL and PP hepatocytes. When a direct comparison was made between our results in these female Sprague-Dawley rats and a study in male C3H/He mice (Braeuning *et al*., 2006), there were similarities in regional expression. *Glul, Oat, Cyp1a2, Cyp2e1, Cyp7a1,* and *Nr1i3* were higher in CV and *Pck1, Cyp2f2/4, Aldh1b1, Hes1,* and *Cdh1* were higher in PP in both species.

### TCDD-treated rats

The previous studies with these rats, intended to assist in risk assessment evaluations with TCDD based on liver cancer in female rats, examined the phenotypic outcomes of TCDD in these wild-type and AHR-knock out rats (Harrill *et al*., 2016), WT-but not the KO rats had dose-dependent decreases in body weight and increases in liver-to-body weight ratios (Harrill *et al*., 2016). Using liver tissues from that study, we found that WT, but not the AHR-KO rats, also displayed dose-related decreases in cell replication in the liver Ki67 labelling (**Figure 2A-B**). This group of tissue responses – loss of body weight, increased liver to body weight ratios and decreased cell proliferation - indicate that the doses evaluated in this study are at or approaching levels associated with wasting and examining DEGs in the livers from these rats should provide information on the metabolic status of the liver at doses causing toxic signs consistent with wasting.

**Figure 2:**
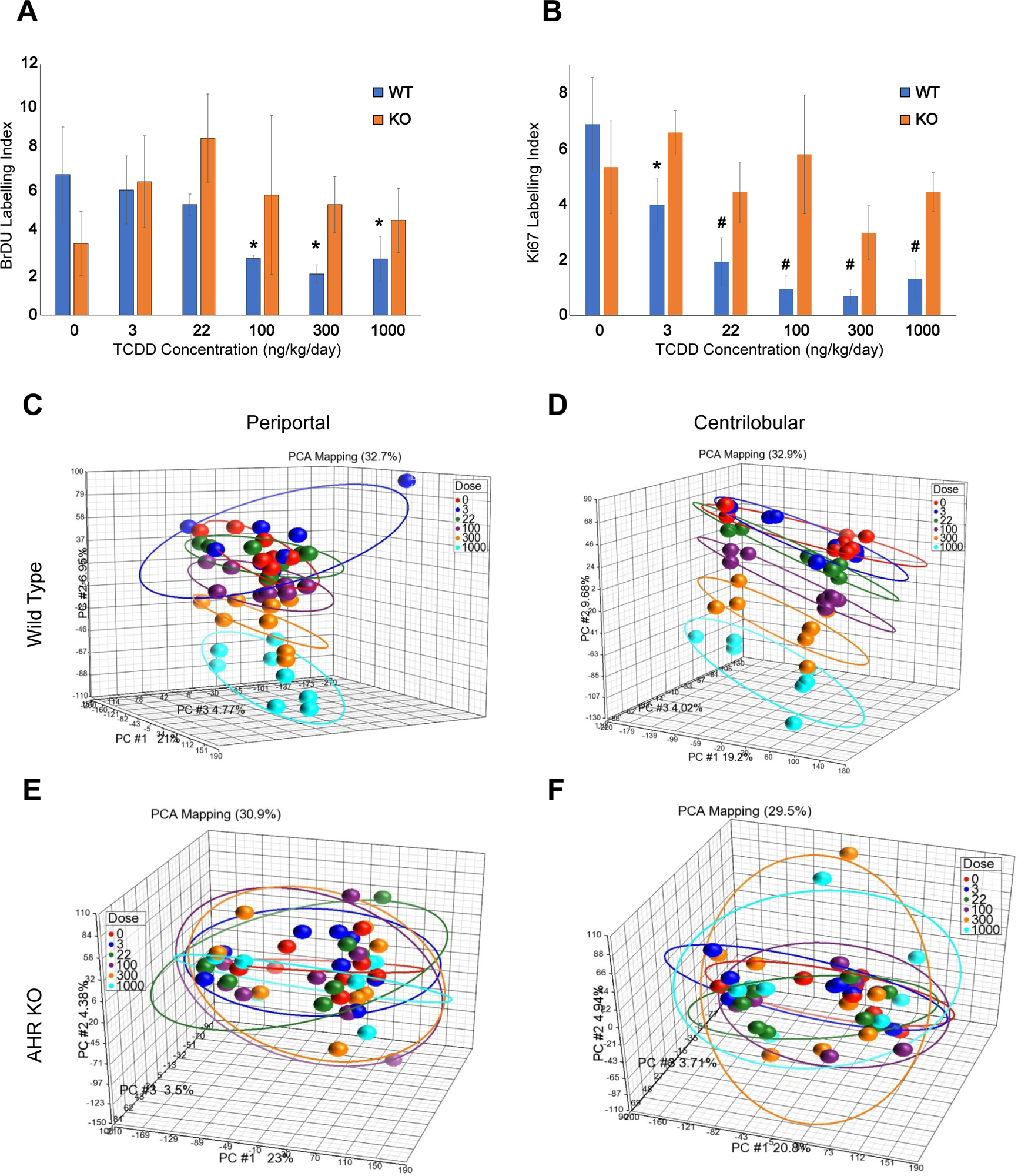
Proliferative indices of wild-type and AHR knockout rat livers in response to TCDD treatment. Rats were administered TCDD via oral gavage daily for 28 days (5days/week). Tissues were collected at the terminal endpoint. TCDD treatment caused a dose-dependent decrease in both BrdU (A) and Ki67 (B) positive cells in WT but not KO rat livers (p<0.05, p<0.001 by ANOVA) and the decrease is generally greater with Ki67. PCA analyses of differentially expressed genes in Wild-type and AhR-KO (C-H) rats after administration of various doses of TCDD. The separation of dose groups is clearest for the CL region (E) and only apparent at higher doses in the PP hepatocytes. There is no clear separation across dose in either liver region with the AhR-KO rats (F-G).

### Differential gene expression

Microarray analysis of gene expression in wildtype rat CL or PP liver regions treated with various doses of TCDD showed dose-dependent increases in the number of differentially expressed genes (DEGs) using criteria of FDR < 0.05, |FC| > 1.5 (**Table 1**). At each dose, there were more DEGs in the CL region of the livers when compared to the PP region and the higher doses there were large increases in the number of both upregulated and downregulated genes. Very few DEGs were found in AhR-KO rats. A principal component analysis (PCA) plot of the gene expression results for individual WT animals showed the separation among doses and the increased sensitivity of the CL region (i.e., there was a more distinct separation between groups at lower doses with WT CL samples than with WT PP samples). No separation across doses was observed in AhR-KO CL or PP samples (**Figure 2 C-F**).

**Table 1.**
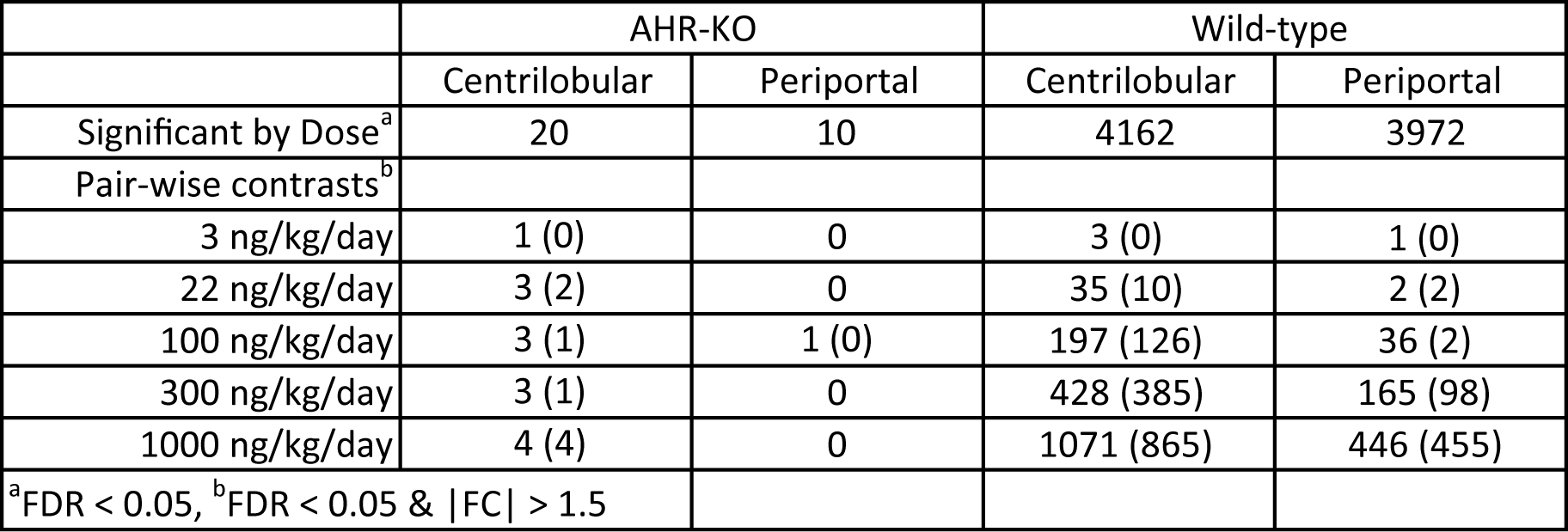
Summary of differential gene expression results with TCDD in the CL and PP regions at different daily doses: number of upregulated genes without parentheses and the number of downregulated genes in parentheses.

In WT samples, there were only 3 upregulated genes at 3 ng/kg/day – *Cyp1a1, Cyp1b1* and *Nqo1* in the CL region. By 22 ng/kg/day in WT samples, 33 of the 45 DEGs were upregulated and included *Cyp1a1, Cyp1b1*, as well as *Cyp1b2, Nqo1* and *Nfe2l2*. In the PP region, at 22 ng/kg/day in WT samples, only two genes were increased (*Cyp1a1* and *Cyp1a2*) while 31 of the 38 DEGs at 100 ng/kg/day were upregulated. These genes included *Cyp1a1, Cyp1a2, Cyp1b1, Nf2el2, Nqo1* and *Ahrr*. This canonical group of xenobiotic metabolizing genes represented the most sensitive (i.e., dose responsive) genes in both liver regions of WT samples while the dose response curves for *Cyp1a1, Cyp1b1, Nqo1 and Ahrr* were also right shifted for the PP compared to the CL region (**Figure 3**). These genes were increased at all doses and no dose-responsive changes in DGE were observed in AhR-KO rats.

**Figure 3:**
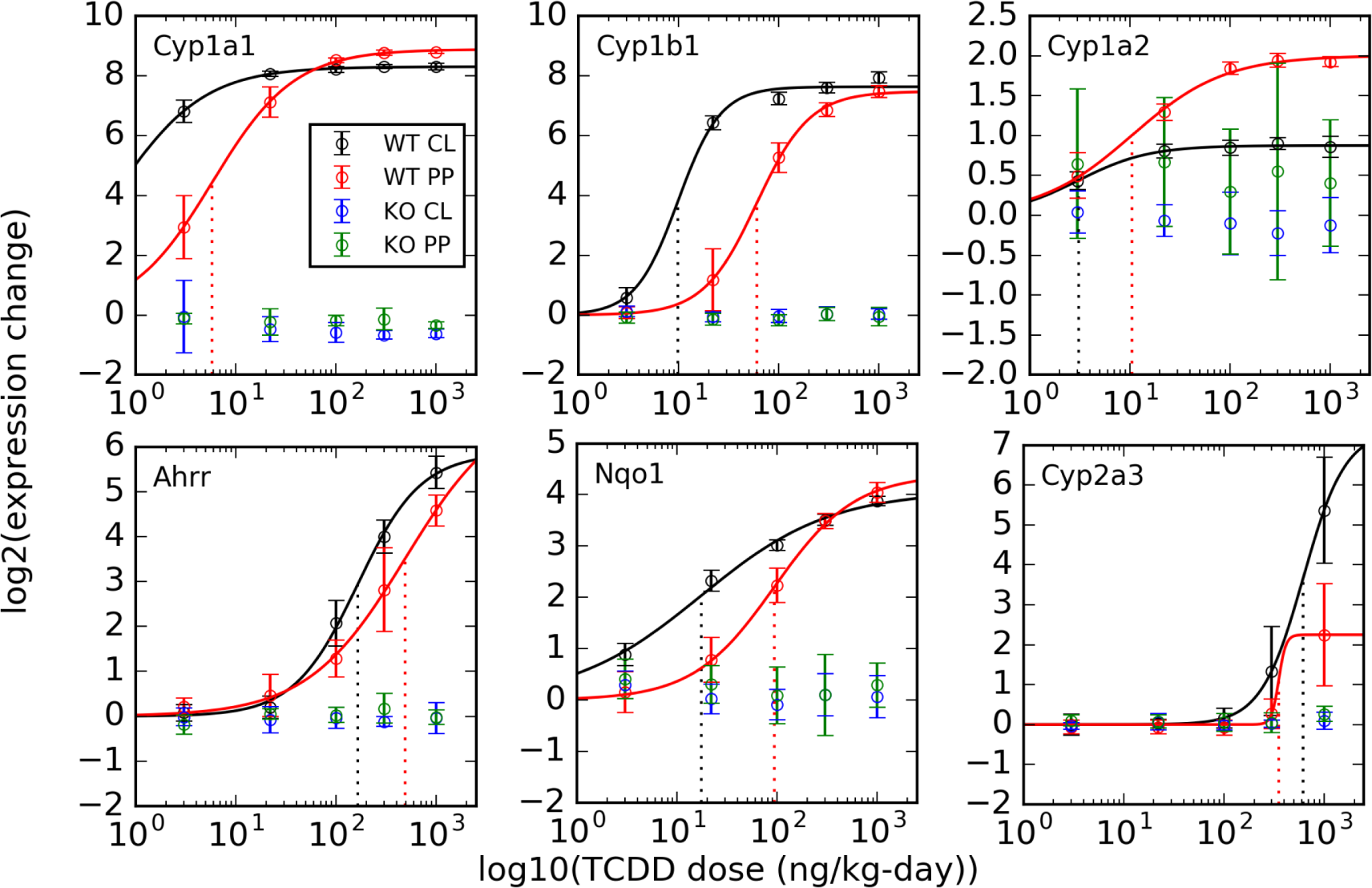
Dose response behavior for selected genes whose activity is controlled by DRE-binding of AHR. Few DEGs were found with the AHR-KO rats. The right shifted dose response curves are most clear with *CYP1a1, Cyp1b1,* and *Nqo1* and the dose-response is left shifted for gene expression in the CL region.

### Transcription Factor Enrichment for DEGs

Although the DEGs in the two WT regions differed, transcription factor binding (TF) enrichment, determined using the Enrichr tool (Xie *et al*., 2021), for the groups of genes at equivalent, low dose response levels (i.e., the 33 upregulated genes in the CL region at 22 ng/kg/day and 31 upregulated genes in the PP at 100 ng/kg/day) were similar and showed enrichment for AHR and ARNT (**Figure 4; Panel C &D**) with a tendency for statistically greater representation of AHR with the CL genes and ARNT with the PP genes.

**Figure 4:**
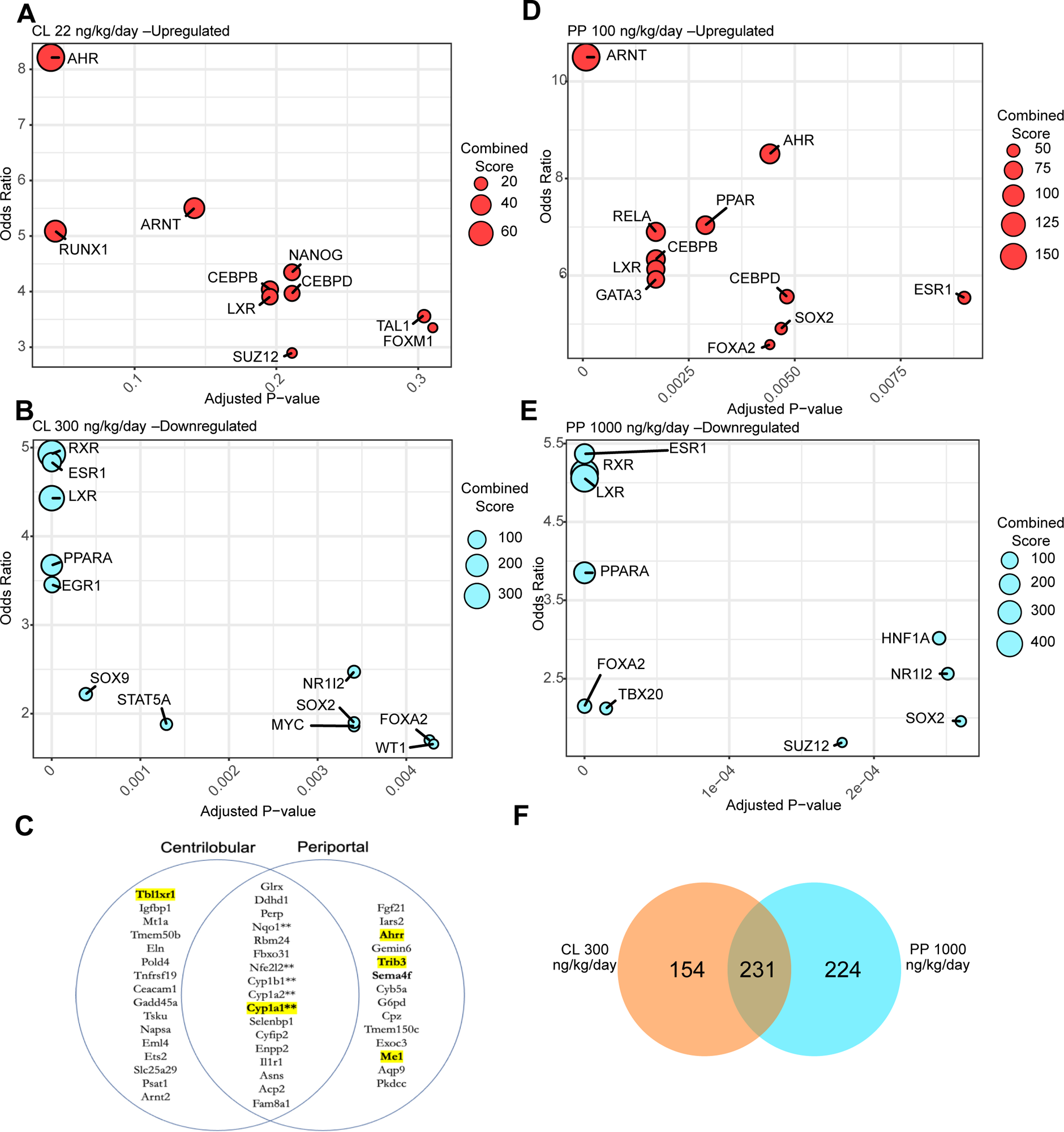
Enrichr TF analysis identifies similar enrichment in H-WT CL and PP hepatocytes with differing doses of TCDD. (A) The overlap of upregulated genes for 22 ng/kg/day TCDD in CL and 100 ng/kg/day in PP hepatocytes. At these lower doses, individual genes are explicitly identified. (B) Overlap of downregulated genes at doses causing similar numbers of genes in CL (300 ng/kg/day) and PP (1000 ng/kg/day) hepatocytes. Panels C and D show Enrichr transcription factor enrichment for upregulated genes for CL and PP hepatocytes, respectively. Panel E and F show Enrichr TF plots for downregulated genes for CL and PP hepatocytes, respectively. The odds ratio is a statistical measure used to assess the enrichment of transcription factor binding sites (TFBSs) within the regulatory regions of genes in a specific gene set compared to a background reference gene set and indicates whether the TFBSs associated with a particular transcription factor is significantly overrepresented or underrepresented.

Similarly, the specific genes in the two liver regions for downregulated DEGs at doses causing approximately equivalent number of DEGs (for instance, 300 ng/kg/day in CL versus 1000 ng/kg/day in PP) also differed, yet TF enrichment profiles were similar (**Figure 4; Panels E and F).** Among the top 10 enriched TFs for these downregulated DEGs at these higher doses were 7 nuclear receptors, RXR, LXR, PPARα, ESR1, NR1I2, FOXA2 and SOX2, common to both liver regions, as well as unique hits, such as EGR1 in CL and HNF1A in PP regions.

With upregulated genes at the 1000 ng/kg/day doses, TF enrichment displayed a more diverse group of TFs with lesser involvement of nuclear hormone receptors (**Supplementary Figure S1**). In the CV region, TF enriched for the upregulated genes included SOX2, CEBPD, TCF2, NRF2 and SMAD 2 and 3; in the PP region, they included NRF2, SMADs 2, 3 and 4, and SOX2. TF binding enrichment for PPAR γ was seen for upregulated DEGs in both regions.

### Visualization of Gene Ontology (GO) pathway analysis

For TF enrichment (Fig 4), the DEGs for specific treatments were transported to a platform (Enrichr) that querues various data bases to evaluate the frequency of TF binding motifs in the gene sequences for each of the DEGs. We also did pathway entichment for the DEGs themselves using the Reactome ontology (Gillespie *et al*., 2022) and CytoScape software to visualize GO-association network graphs for DGE identified for each condition (McMullen *et al*., 2019). Visualization of these network graphs for up and down regulated genes for 1000 ng/kg/day in CL show upregulation of pathways for extracellular matrix organization, hemostasis and innate immune system (**Figure 5**) with extensive downregulation of genes in pathways for metabolism of fatty acids, ketone body metabolism, metabolism steroids, pyruvate metabolism and citric acid (TCA) cycle, biological oxidations, metabolism of amino acids and derivatives, metabolism of vitamins and co-factors and gluconeogenesis. This finding is consistent with the report that rats treated with doses of TCDD sufficient to cause signs of wasting are not protected by total parenteral nutrition (Gasiewicz *et al*., 1980). The upregulation of pathways for innate immune system, hemostasis and extracellular matrix organization were found at the highest dose in the CL region.

**Figure 5:**
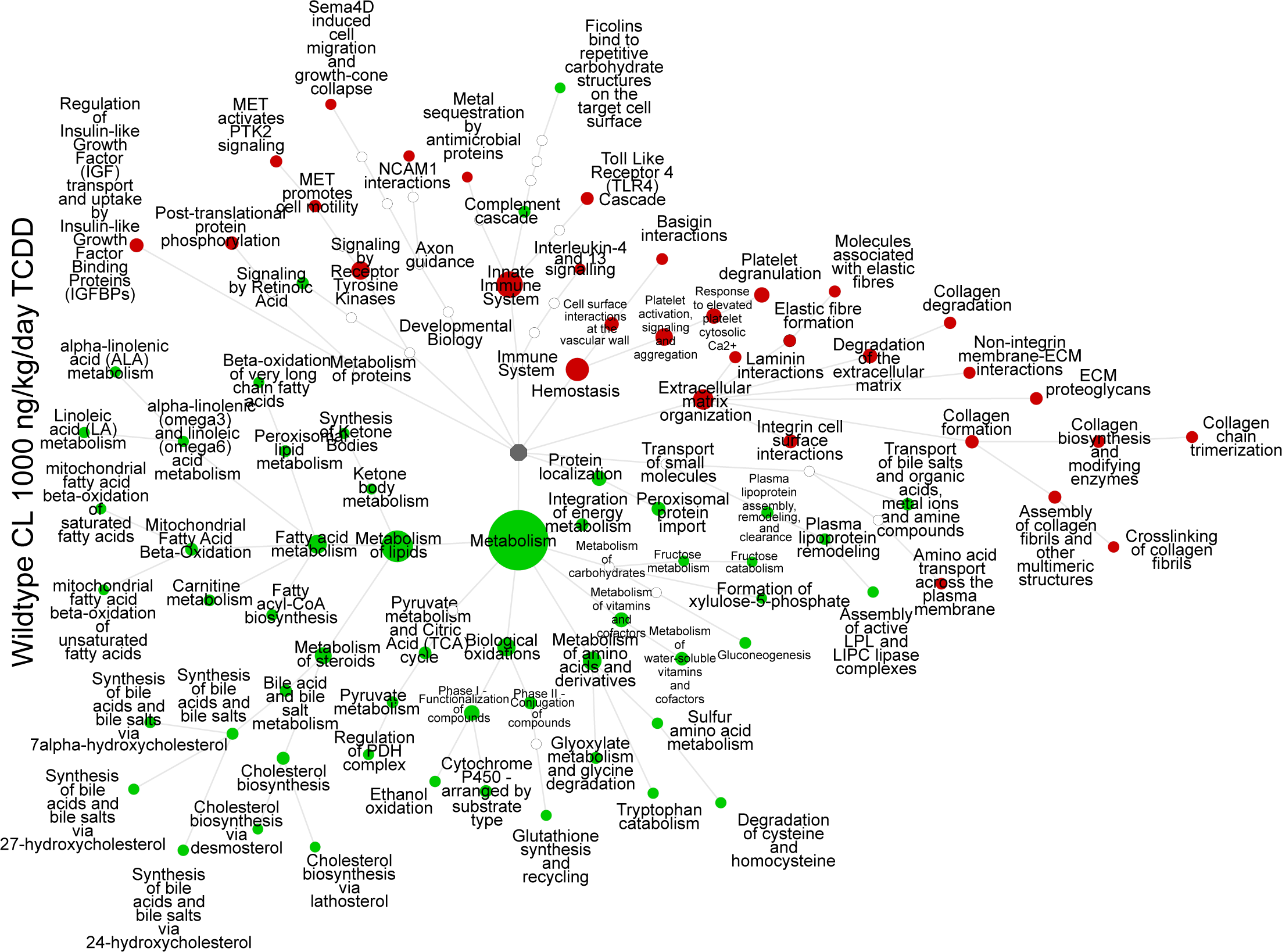
Pathway visualization maps of Reaction pathway enrichment for up and downregulated pathways in CL hepatocytes following 1000 ng TCDD/kg/day. There are 44 genes in fatty acid metabolism pathway and 34 genes in the metabolism of steroids pathway and a variety of upregulated pathways associated with innate immune system and extracellular matrix organization. In all these GO-category visualization plots the size of the circles correspond to the total number of genes in the pathway. Green represents downregulated pathways; red, upregulated pathways.

The downregulated pathway that was most significantly affected in the CL region at 1000 ng/kg/day was activation of gene expression by SREBP with 34 genes, 23 of which are in cholesterol biosynthesis pathway. In addition to downregulation of both *Srebp1* and *Srebp2*, affected genes for cholesterol biosynthesis included *Hmgcs1* (hydroxymethyl-glutaryl CoA synthetase), *Tmf7sf2* (reduction of lanosterol), *Cyp51* (lanosterol demethylation), *Lss* (lanosterol synthetase), *Sc5d* (sterol-5 desaturase), *Pmvk* (phosphomevalonate synthetase), *Fdps* (farnesyl diphosphate synthetase), *Idi1* (isopentenyl-diphosphate decarboxylase) and *Mvd* (mevalonate diphosphate decarboxylase). Among the 44 downregulated genes in fatty acid metabolism were *Acaca* (acetyl CoA carboxylase), *Fasn* (fatty acid synthetase) and *Gpam* (glycerol-3-phosphate acyl transferase) (**Figure 5**). In comparing the 1000 ng/kg/day PP region response to that at 300 ng/kg/day in the CL, the plots were similar although a larger number of genes were altered in each of the common pathways in the PP region (**Figure 6**). At doses causing clear wasting-like responses with TCDD in these rats (300 to 1000 ng/kg/day), there was significant downregulation of pathways for fatty acid metabolism, metabolism of steroids, biological oxidations, metabolism of amino acids and derivatives, and pyruvate metabolism and citric acid cycle only in the CL hepatocytes at 1000 ng/kg/day (Table 2). At 100 ng/kg/day in the CV hepatocytes and 300 ng/kg/day in the PP hepatocytes, there was downregulation of the general pathway of metabolism represented by 38 and 33 genes respectively. However, the genes represented multiple different pathways without enrichment of fatty acid metabolism or metabolism of steroids. The core set of TCDD responsive genes, *Cyp1a1, Cyp1a2, Cyb1b1 and Nqo1* were upregulated at all TCDD doses (**Table 2**).

**Figure 6:**
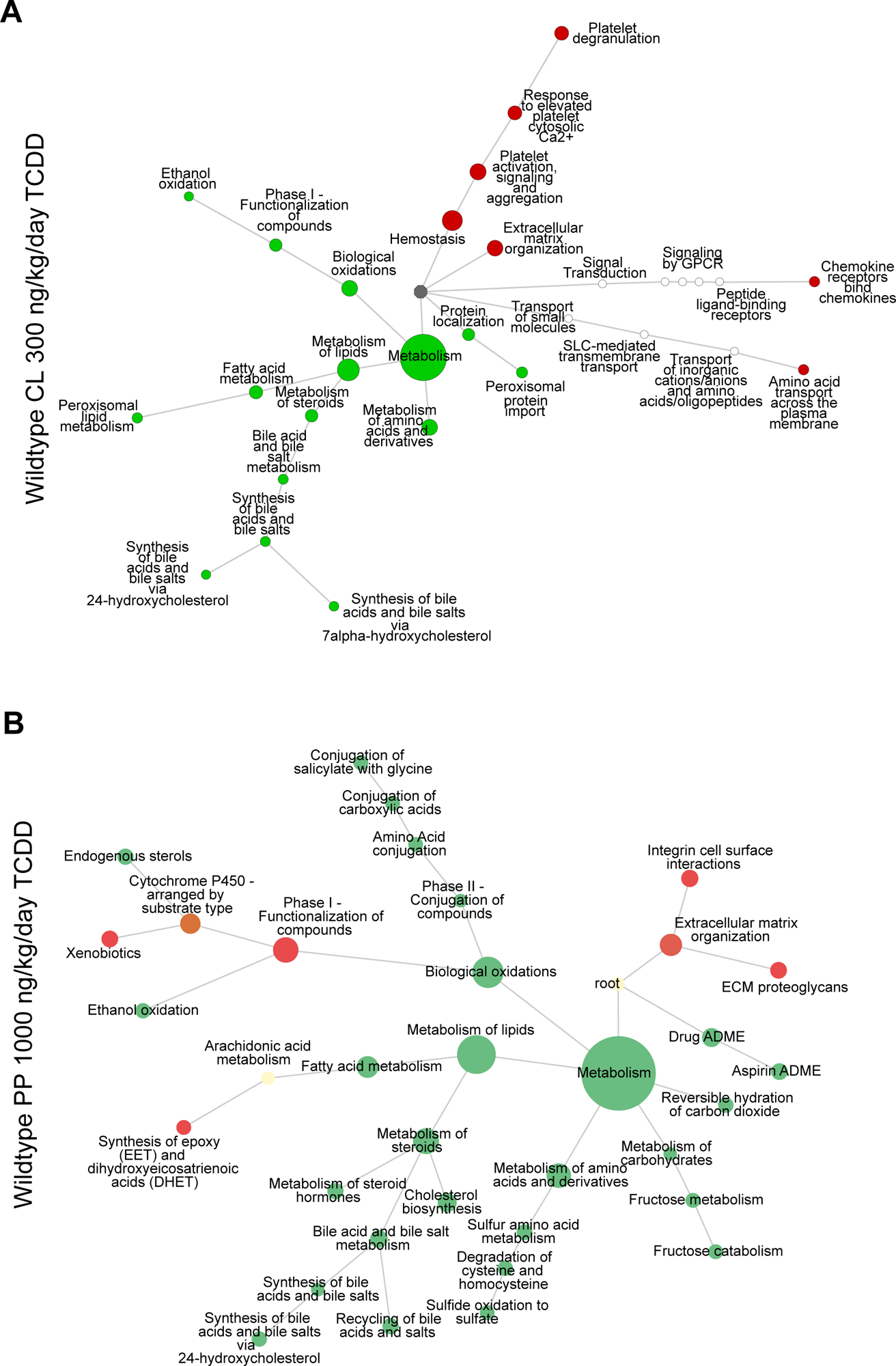
Pathway visualization maps of Reactome pathway enrichment for up and downregulated pathways in CL hepatocytes following A) 300 ng TCDD/kg/day compared to those in the B) PP region at 1000 ng/kg/day. At this dose in the CL region, there were 15 genes in the fatty acid metabolism pathway and 13 genes in the metabolism of steroids pathway with much more limited enrichment for upregulated pathways of hemostasis and extracellular matrix organization. In PP hepatocytes, a larger number of pathways were downregulated and there were more genes in the pathways common: 18 in fatty acid metabolism and 27 in metabolism of steroids.

### Differential Expression of Transcription Factor Genes

At 300 and 1000 ng/kg/day there was enrichment with downregulated genes for various metabolic processes and for pathways of extracellular matrix, hemostasis, and innate immune system with upregulated genes (**Figure 5**). At these higher doses, a variety of TF genes were also differentially regulated (**Fig S2).** In the CL region at 1000 ng/kg/day there were 62 upregulated and 34 downregulated TF genes. At the same dose in the PP region there were 29 up- and 22 down-regulated TF genes. Among the upregulated nuclear hormone receptor subfamily genes at 1000 ng/kg/day in both liver regions were *Nr1d1, Nr1d2,* and *Nr4a1. Nr1i3*. *Ppara* was only increased in the PP region and *Nr5a2* in the CL region. Among the downregulated nuclear receptor hormone subfamily TF genes in the CL region were *Nr1h4, Nr3c1, Rarb, Parg and Ppard* and in the PP region the NR family genes downregulated were *Esr1, Nr1h4, and Ppard*.

Several *Bhlhe* TF genes we also affected. At 1000 ng/kg/day, *Hes1* and *Heyl* were upregulated in the CV region and *Mlxipl, Bhlhe40*, *Arntl*, *Srebf1*, and *Srebf2* were downregulated. In the PP region at this dose, *Ahr* was upregulated and *Srebf1, Mlxipl, Bhlhe40* and *Hes6* were downregulated. HES1, HES6, HEYL and BHLHE40 are transcriptional repressors (Kageyama *et al*., 2000; Gao *et al*., 2001; Katoh and Katoh, 2007; Nakashima *et al*., 2008; Han *et al*., 2020). MLXIPL, SREBF1 and SREBF2 positively regulate the synthesis of cholesterol, lipids and fatty acids (Song *et al*., 2018). Protein products of two other TF genes, *Hnf1a* and *Hnf6a*, are involved in regulating liver specific gene expression, were both downregulated in the CV region and, of these two, *Hnf1a* alone was downregulated in the PP region. These genes play important roles in liver development (Tachmatzidi *et al*., 2021).

## DISCUSSION

### LCM and Regional Differences in Liver Responses to TCDD

LCM gave two populations of hepatocytes with differences in gene expression in vehicle control rats consistent with other reports (Braeuning *et al*., 2006). There were also different dose-response behaviors with TCDD for the two regions with the PP tissues being less sensitive (i.e., right shifted compared to the CV region). Nonetheless, the low dose responses in each area were consistent with activation of AHR and ARNT signaling. The greater sensitivity of the CV region may be a consequence of a functional role of the liver in clearing biologically active compounds from the circulation where blood exits the liver to return to the general circulation. As concentrations increase throughout the body induction may spread to the PP region to increase effective clearance. Alternatively, the CL-hepatocyte TCDD concentrations were found to be 2.7 to 4.5-fold higher than those in the PP (Santostefano *et al*., 1999) and these concentrations differences may drive the different dose response between regions. The differences in regional TCDD are likely due to differences in regional CYP1A2 levels, a protein to which TCDD is strongly bound (Poland *et al*., 1989).

### Wasting and Altered Metabolism

Under conditions leading to wasting with TCDD, gene expression in liver showed downregulation of many pathways for metabolism. At 300 ng/kg/day, Ki67 staining, that labels cells at the time of collection, showed nearly complete cessation of cell proliferation (Figure 2). The differences in Ki67 and lesser response with BrdU staining are consistent if longer exposure times are required to alter proliferation. BrdU integrates proliferation over a 5-day period at the end of the study; Ki67 stains tissues undergoing proliferation at the time of necropsy. With respect to the dose response for cell proliferation (Figure 2), Ki67 is affected at lower doses and the decrease compared to control is larger than with BrdU. Overall, the dose response for decreased proliferation is more consistent with the dose response for increases in the DRE-containing cytochrome genes, *Cyp1a1, Cyp1a2* and *Cyp1b1* (Figure 3). The dose response for downregulation of lipogenic genes that lack DREs is shifted to higher doses than that for activation of canonical AHR responsive genes and that the upregulation of genes involved in extracellular matrix, innate immune system and hemostasis appears to shift to even higher doses.

Down regulation of cholesterol biosynthesis pathways by TCDD had been previously noted in mice (Sato *et al*., 2008; Tanos *et al*., 2012). Here we have shown that (1) similar effects occur in rats; (2) the dose response for these effects coincides with that for wasting; and (3) these responses are absent in AhR-KO rats. Changes in cholesterol pathway genes in mice were also independent of the presence of AHR-dioxin response element. The responses occurred in mice with an AHR-DRE binding mutation - *AhR-A78D* (Tanos et al., 2012). It is now clear that TCDD at higher doses (consistent with wasting type responses or near wasting doses) affects a much larger suite of pathways of metabolism (**Figure 5**) and also upregulates genes for pathways for extracellular matrix (ECM), hemostasis and innate immune system. While our studies did not include determination of protein levels for differentially expressed genes, the changes in pathway genes expression batteries of genes in lipogenesis and cholesterol synthesis emphasizing the coordinated, sequential alteration in these pathways.

With TCDD, the lower doses (22 ng/kg/day in CL and 100 ng/kg/day in PP) upregulated genes controlled by the AHR and ARNT – among them the three *Cyp* genes and *Nqo1*-without downregulation of various pathways of cellar metabolism. Only at 300 ng/kg and above in the CL region was there evidence for down regulation of multiple pathways of cellular metabolism and at the very highest doses evidence for upregulation of pathways associated with ECM organization, hemostasis and innate immune system. A strong case can be made that wasting, at least at the level of altered metabolic processes, with TCDD is primarily associated with inhibition of lipogenesis, other pathways of cellular metabolism and cell proliferation, and subsequent activation of pathways for extracellular matrix organization. The present study strongly indicate that prolonged inhibition of various anabolic pathways and upregulation of ECM pathways lead to wasting and that responses at these higher dose responses are not due AHR binding to DREs.

In addition to the extensive downregulation of TF genes in pathways for metabolism, there was downregulation of two key TF genes that control liver development and adult liver gene expression, *Hnf1a* and *Hnf6a* (Onecut1) (Tachmatzidi *et al*., 2021). Proteins transcribed from these genes are involved in maintaining expression of a variety of liver transcripts. The net results at the higher daily doses of TCDD is downregulation of liver associated metabolic pathways, upregulation of genes in pathways for extracellular matrix, hemostasis and innate immune system. Overall, TCDD treatment showed three distinct regions with different responses. At the lower doses, we found canonical expression of DRE-responsive genes (Figure 4A and D) at the intermediate dose near 300 ng/kg/day, there was decreased expression of genes for multiple metabolic pathways. At the highest dose, a wider array of upregulated pathways became prominent. Overall, the biology of liver responses to TCDD range through processes that include a normal response to the presence of excess biologically active compound proceeding to changes in gene and TF expression suggestive of dedifferentiation of hepatocytes and finally to gene and TF factor changes suggestive of epithelial-mesenchymal transition (Kalluri and Weinberg, 2009).

### Possible molecular targets leading to altered lipid metabolism and wasting by TCDD

The negative feedback control is an important component of circadian rhythms (see Supplementary Discussion Material) and feedback of NR1D1 and NR1D2 on regulation of circadian controlled genes. The CNS clock is driven by changing patterns of daylight. Light activates pathways in the super-chiasmatic nucleus (SCN) affecting other brain regions. Signals for light-period responses in peripheral tissues are not yet well understood but may be driven by production of high affinity AhR-active ligands, such as the tryptophan photodimer, FICZ (Rannug and Rannug, 2018) or by feeding. FICZ binds AHR, leading to its transport from the cytoplasm to the nucleus with formation of the AHR-TCDD-ARNT complex. Dimerization of AHR and ARNT shifts their activity to control of Phase I and Phase 2 genes coding for proteins that accelerate clearance of small molecules (i.e., the gene expression changes occurring at the lower doses in this study). Even at the lower doses with TCDD, there was upregulation of *Cyp1a1, Cyp1a2* and *Cyp1b1* which remain maximally elevated as TCDD dose increased (**Figure 3**). At the end of the light cycle, melatonin is released from the pineal gland into the general circulation. The exact role of melatonin in the circadian control of metabolism in peripheral tissues remains uncertain. This hormone interacts with two G-protein coupled melatonin receptors – MTNR1A and MTNR1B - activating AMP kinase - AMPK (Rui *et al*., 2016), a process that could target CRY for degradation. Melatonin is metabolized to 6-hydroxymelatonin by *Cyp1a1, Cyp1a2* and *Cyp1b1* (Ma *et al*., 2005) and its clearance from circulation would be increased by high dose TCDD exposures with persistent increases in these cytochromes (Fig 3). Single intraperitoneal doses of TCDD (50 ug/kg) caused prolonged, significant reduction in circulating melatonin (Linden *et al*., 1991; Pohjanvirta *et al*., 1996).

### A Model for TCDD effects on Circadian Clock and AhR in circadian control

Our DGE results support a hypothesis for circadian control in peripheral tissues where effects of TCDD on gene expression would not depend on direct binding of AHR-ARNT to DREs. In this hypothesis (Figure 7), AHR and ARNT serve as integrators of the multiphasic control of circadian cycling by (1) responding to light induced ligands; (2) inducing metabolism of excessive concentrations of ligands that are capable of binding AHR; (3) taking part in fundamental biology of repair and regeneration in the rest phase of the circadian cycle; (4) modulating gene expression behaviors controlled by BMAL1-CLOCK; and, (5) serving to enhance clearance of melatonin at the beginning of the light phase. Our results suggest that persistence of high tissue exposures of TCDD, a high-affinity, poorly cleared AHR agonist, could stall the circadian cycle, sequester AHR and ARNT and maintain high levels of cytochromes that would metabolize melatonin, FICZ or other molecules involved in the shift of metabolism from the activity to the rest phases of the circadian cycle. TCDD also would titrate AHR away from its permissive role in cell proliferation, a response that could be responsible for the reduced cell proliferation across most of the doses seen with Ki67 labeling (Figure 2) and titrate ARNT away from HIF-1A activity as a hypoxia (i.e., low oxygen) responsive dimer.

**Figure 7:**
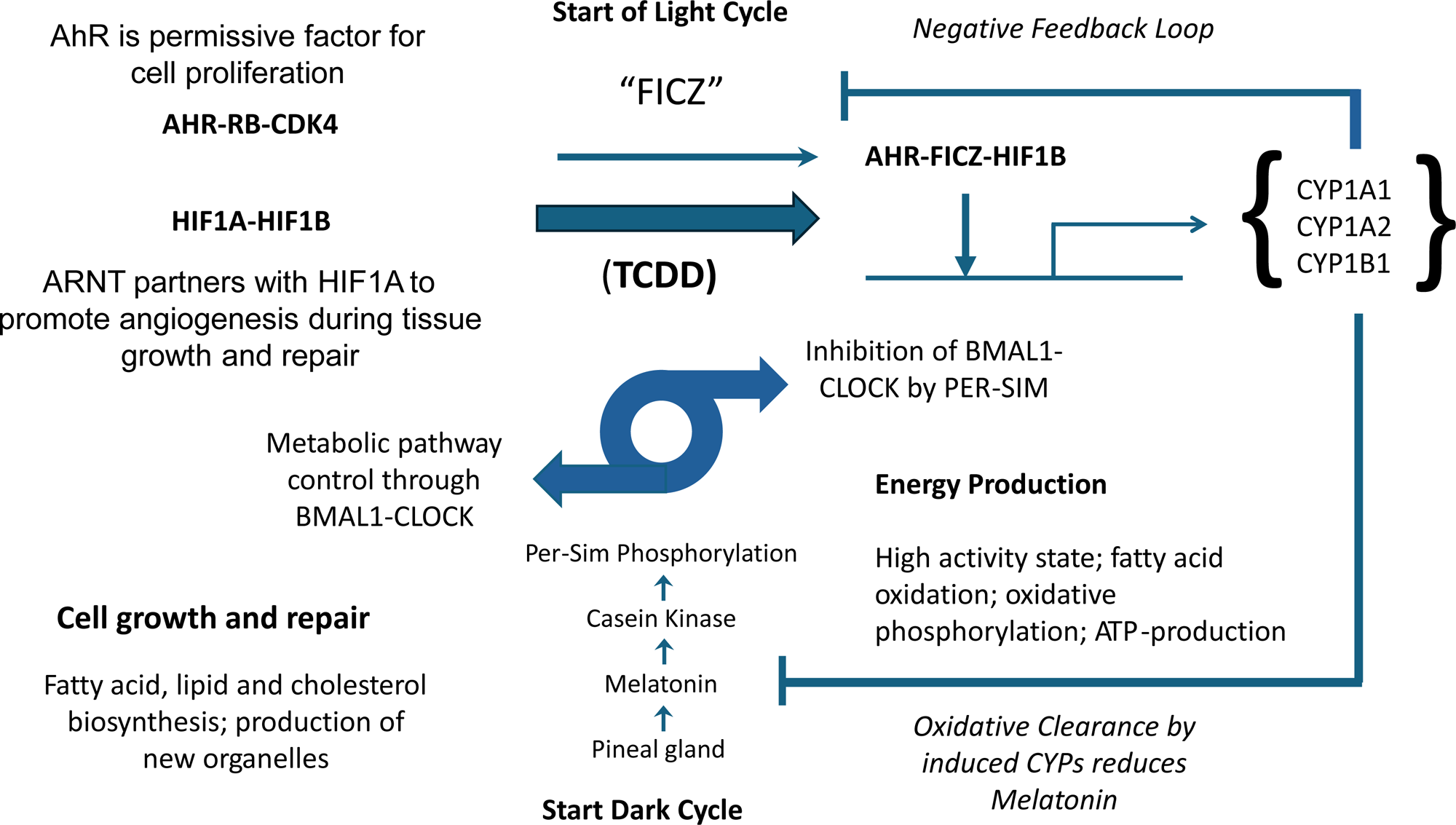
Proposed model for integration of AHR and ARNT in circadian cycling and effects of TCDD. Starting at the end of the dark cycle where BMAL1 and CLOCK would regulate metabolic output, light produces AHR-active ligands. The activation by ligand would titrate both AHR and ARNT from non-liganded forms and increase enzymes that clear the bioactive ligand and melatonin. As light intensity diminishes later in the light cycle, melatonin released from the pineal gland, would circulate systemically and, through activating melatonin receptors and kinases, AMP-K and phosphorylate of PER and CRY promoting their degradation and diminishing their inhibition of BMAL1-CLOCK. The persistence of a bioactive, poorly metabolized ligand, TCDD, at high levels would stall the circadian cycling of gene expression, inhibit permissive roles in cell proliferation (AHR) and angiogenesis (ARNT), increase the clearance of melatonin and cause downregulation of multiple metabolic pathways and ultimately wasting.

A companion paper examines *in vitro* DGE in human hepatocytes treated with TCDD for up to 14-days. Several PFAS – specifically perfluorooctane sulfonic acid, perfluorodedecane sulfonic acid and perfluoro-n-decanoic acid – show profound downregulation of lipogenesis and upregulation of celllar stress pathways (Barutcu *et al*., 2024). The changes in lipogenesis are a common signal with wasting/failure to thrive responses with these diverse compounds while the MOA for TCDD and these longer chain PFAS likely varies substantially.

## Supporting information

Supplementary Discussion

## Acknowledgements

The *in vivo* studies with TCDD and collection of gene expression data from livers of the rats in the study were supported by a grant from the Dow Chemical Company to the Hamner Institutes. The bioinformatic and computational tools for analyzing gene expression, pathway enrichment and network visualization were subsequently developed with funding provided to the Hamner and Institutes and ScitoVation LLC by the Long-Range Research Initiative of the American Chemistry Council (ACC-LRI). The senior author (MEA) thanks Dr. William F. Greenlee for many fruitful discussions regarding the biological roles of unliganded AHR and ARNT and discussions with Drs. Sudin Bhattacharya and Daniel Mari at Michigan State University for helpful discussions regarding regulation of circadian cycle genes by TCDD. The authors also thank Bethany Parks for expert technical expertise in collection and processing of the LCM zonal hepatocyte samples for gene expression analysis.

## Data Access

The gene expression results have been deposited in the National Center for Biotechnology Information (NCBI) Gene Expression Omnibus (Accession no: GSE72506).

## Supplementary Material

Supplementary Figure S1: TF factor enrichment for upregulated DEGs at 1000 ng/kg/day in both the CV and PP regions of the liver. DEGs were submitted to the Enrichr website (https://maayanlab.cloud/Enrichr) and analyzed using EnrichR 2022.

Supplemental Discussion Material: Negative feedback and NR1D1 and NR1D2 interactions in circadian circuitry.

