## Supplementary figures and images for "Investigating the mode of action for wasting produced by tetrachlorodibenzo-p-dioxin (TCDD) in rats using transcriptomics: Evidence for roles of AHR and ARNT in circadian cycling"

### Supplementary Figure S1

#### Supplementary Figure S1

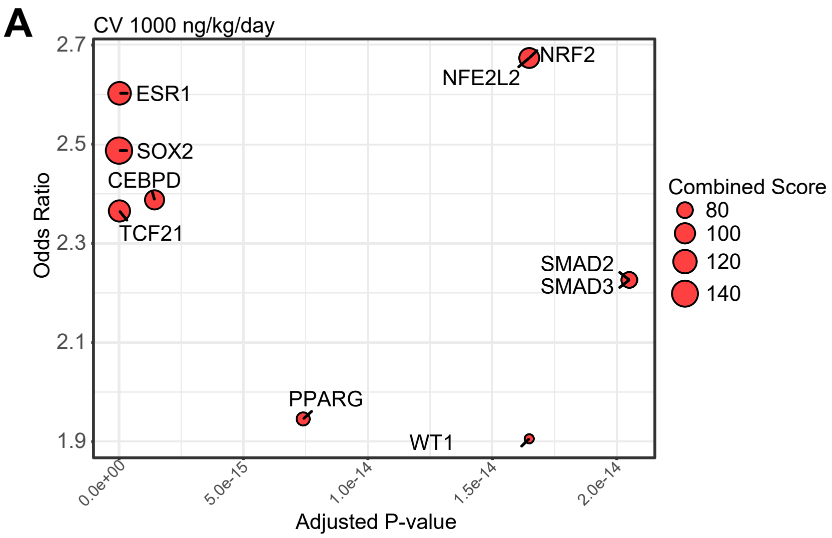


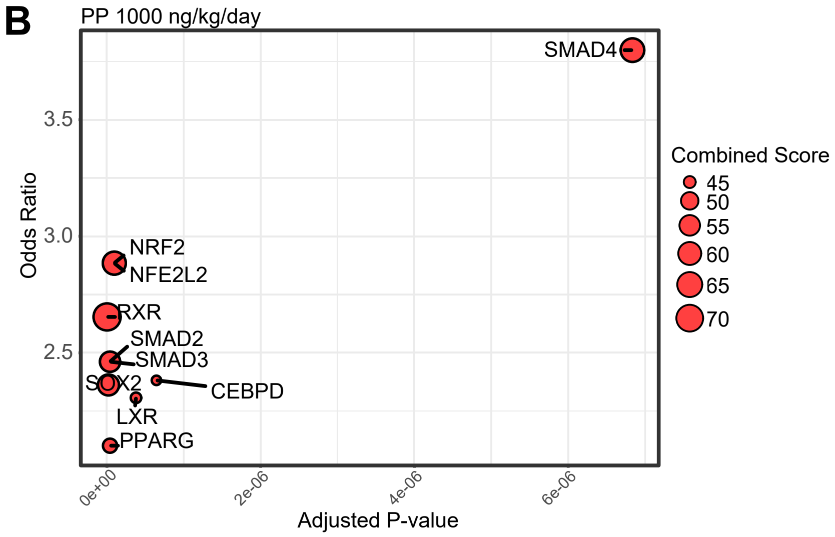
