## Supplementary Discussion for "Investigating the mode of action for wasting produced by tetrachlorodibenzo-p-dioxin (TCDD) in rats using transcriptomics: Evidence for roles of AHR and ARNT in circadian cycling"

**Andersen et al.**

**Supplementary Discussion:**

**Core Elements of Circadian Control of Metabolism**

The circadian clock has a core negative feedback loop (Kohsaka and Bass, 2007; Takahashi, 2017). A dimer of two b-HLH proteins, CLOCK and BMAL1, binds to E-box promoter regions driving expression of direct outputs of the clock and increasing expression of metabolic target genes. The direct outputs of CLOCK-BMAL1 include oscillating RNAs within metabolic networks that control gluconeogenesis, mitochondrial biogenesis, oxidative phosphorylation, amino acid turnover, lipogenesis and bile acid synthesis (Bass and Takahashi, 2010), i.e., many of the same pathways that are affected at the intermediate and highest doses of TCDD. The negative feedback arises from two genes – Period (PER) and Cryptochrome (CRY) - that also are regulated by the BMAL1-CLOCK dimer and form a dimer that inhibits BMAL1-CLOCK. Phosphorylation of PER by casein kinases and CRY by 5'-AMP-activated kinase increases their degradation and relieves the inhibition of CLOCK-BMALs (Kohsaka and Bass, 2007; Takahashi, 2017). Both NR1D1 and NR1D2 repress expression of BMAL1 (Bass and Takahashi, 2010) and alter normal lipid homeostatic gene networks linked to circadian behavior (Cho *et al.*, 2012). While the circadian feedback loops have the capacity to be self-sustaining, they are entrained in the CNS by light-signals through actions of the suprachiasmatic nucleus. The nature of entrainment for systemic tissues is not yet understood, both feeding light cycle behavior and melatonin release at the beginning of the dark cycle may be important for shifting between resting and activity phases of the circadian cycle.
